# Cryo-EM structure of OSCA1.2 from *Oryza sativa:* Mechanical basis of hyperosmolality-gating in plants

**DOI:** 10.1101/505453

**Authors:** Koustav Maity, John Heumann, Aaron P. McGrath, Noah J Kopcho, Po-Kai Hsu, Chang-Wook Lee, James H. Mapes, Denisse Garza, Srinivasan Krishnan, Garry P. Morgan, Kevin J. Hendargo, Thomas Klose, Steven D. Rees, Arturo Medrano-Soto, Milton H. Saier, Miguel Piñeros, Elizabeth A. Komives, Julian I. Schroeder, Geoffrey Chang, Michael H. B. Stowell

## Abstract

Sensing and responding to environmental water deficiency and osmotic stresses is essential for the growth, development and survival of plants. Recently, an osmolality-sensing ion channel called OSCA1 was discovered that functions in sensing hyperosmolality in *Arabidopsis*. Here, we report the cryo-EM structure and function of an ion channel from rice *(Oryza sativa;* OsOSCA1.2), showing how it mediates hyperosmolality sensing and ion permeability. The structure reveals a dimer; the molecular architecture of each subunit consists of eleven transmembrane helices and a cytosolic soluble domain that has homology to RNA recognition proteins. The transmembrane domain is structurally related to the TMEM16 family of calcium dependent ion channels and scramblases. The cytosolic soluble domain possesses a distinct structural feature in the form of extended intracellular helical arms that is parallel to the plasma membrane. These helical arms are well positioned to sense lateral tension on the inner leaflet of the lipid bilayer caused by changes in turgor pressure. Computational dynamic analysis suggests how this domain couples to the transmembrane portion of the molecule to open the channel. Hydrogen-deuterium exchange mass spectrometry (HDXMS) experimentally confirms the conformational dynamics of these coupled domains. The structure provides a framework to understand the structural basis of hyperosmolality sensing in an important crop plant, extends our knowledge of the anoctamin superfamily important for plants and fungi, and provides a structural mechanism for translating membrane stress to ion transport regulation.

## Introduction

Hyperosmolarity and osmotic stress are among the first physiological responses to changes in salinity and drought. Hyperosmolality triggers increases in cytosolic free Ca^2+^ concentration and thereby initiates an osmotic stress-induced signal transduction cascade in plants (1–3). Salinity and drought stress trigger diverse protective mechanisms in plants enabling enhanced drought tolerance and reduction of water loss in leaves.

Ion channels have long been hypothesized as sensors of osmotic stress. A candidate membrane protein named OSCA was isolated in a genetic screen for mutants that impair the rapid osmotic stress-induced Ca^2+^ elevation in plants (1). OSCA1 encodes a multi-spanning membrane protein that functions in osmotic/mechanical stress-induced activation of ion currents. However, the underlying mechanisms and whether OSCA1 itself encodes an ion conducting pore specific for Ca^2+^ requires further analysis. OSCA1 is a member of a larger gene family in Arabidopsis with 15 members (4), and with many homologs encoded in other plants and fungal genomes. Furthermore, evolutionary analyses have revealed that OSCA is distantly related to the anoctamin superfamily, that includes the TMEM16 family of calcium dependent ion channels (5).

## Results

As we were interested to determine whether and how osmolality caused OSCA proteins to respond to osmotic stress in crop plants, we screened five such ion channels from rice, over-expressing them as TEV protease cleavable eGFP fusions in *Pichia pastoris*. The *Oryza sativa* hyperosmolality-gated protein (annotated as *OsOSCA1.2*, GenBank KJ920372.1) was found to have both high levels of protein expression and desirable properties during purification and was therefore chosen for further characterization. We purified the membrane protein to homogeneity (**Fig. S1A**) and determined the oligomeric state of purified OSCA using size-exclusion chromatography coupled to multi-angle laser light scattering (SEC-MALLS) analysis, revealing the detergent solubilized protein to be a dimer (**Fig. S1B**).

### Functional Reconstitution of OsOscA1.2

Reconstitution of the OsOSCA1.2 purified proteins into droplet interface bilayers (DIBs) indicated the purified protein is fully functional, mediating ion transport (**Fig. S1C-H**). OsOSCA1.2 was active in symmetrical (*cis:trans* 150:150 mM KCl) ionic conditions in the absence of any other osmotically active solutes (**Fig. S1C-E**). In symmetric ionic conditions the current to voltage relation for OsOSCA1.2 was quite linear, yielding a unitary conductance of 284± 2 pS and showing no signs of current rectification. The unitary conductance is within the range of those reported recently for other OSCA proteins (e.g., between 300 to 350 pS in similar ionic conditions (4). Under non-symmetrical ionic conditions (*cis:trans* 15:150 mM KCl), with an inwardly directed K+ gradient, the inward single channel currents reversed (E_rev_) at about −26mV, closer to the Nernst potential for K^+^ (E_K+_: 54 mV after correction for ionic activities), indicating a modest selectivity for K^+^ over Cl^-^ as suggested by the calculated P_K_+/P_C^-^_ = 5 ± 1 (**Fig. S1I**). The appearance of infrequent 50% current amplitude sub-conductance state (**Fig. S1H**) was consistent with the proposed assembly of two cooperative subunits as confirmed by the dimeric nature of the OsOSCA1.2 channels inferred from SEC-MALLS (**Fig. S1B**). The result is also consistent with and other recent studies of OSCA proteins (6, 7). Overall, the above experiments confirmed functionality of our purified OsOSCA1.2 protein.

### Structure of OsOSCA1.2

We determined a molecular structure of OsOSCA1.2 by single-particle cryo-electron microscopy to an overall resolution of 4.9 Å and local resolution in the membrane of 4.5 Å, revealing a dimer of C2 symmetry related subunits (**Fig. S2**). The overall dimension of the protein are 140 Å x 55 Å x 85 Å. Each protomer is comprised of eleven transmembrane (TM) spanning segments, associated extra- and intracellular loops and an intracellular soluble domain (**Fig. 1A**). All eleven transmembrane helices and the soluble domain are well resolved in our cryo-EM maps and large side chains provided suitable markers for ensuring proper sequence registration during atomic model building. (**Fig. S3**). The final atomic model comprises 1388 out of the expected 1424 residues with good geometry and an EMringer (8) score of 0.89 (**Table S1**).

**Figure 1.**
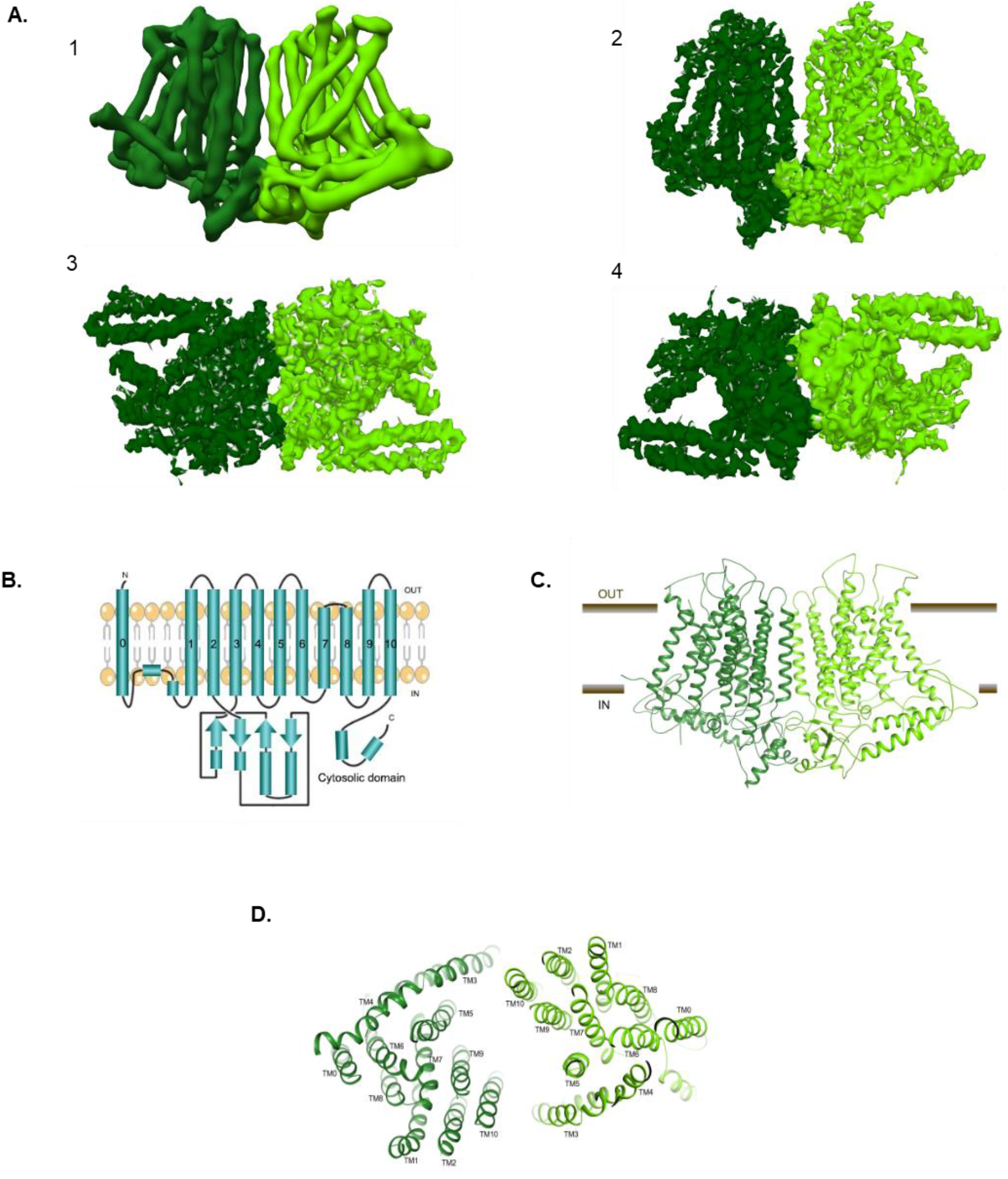
Cryo-EM structure of the OsOSCA1.2 ion channel. (A) From left to right, (1) Parallel to membrane plane view of unsharpened cryoEM density map used for initial chain tracing, (2–4) sharpened 4.9 Å map used for model building and refinement: (2) membrane plane view, (3) extracellular view, and (4) intracellular view. (B) Protein topology of OsOSCA1.2. View of OsOSCA1.2 model from (C) the plane of the cell membrane and (D) from the extracellular side.

According to the Transporter Classification Database (TCDB) (9), OsOSCA1.2 belongs to what is annotated as the Calcium-permeable Stress-gated Cation Channel family (CSC; TC: 1.A.17.5) within the Anoctamin Superfamily (TC: 1.A.17). This classification indicates that OsOSCA1.2 is distantly related to members of the Anoctamin family (ANO; TC: 1.A.17.1) for which high-resolution 3D structure are available (10, 11). Following a recently published bioinformatics approach (5), we had further predicted that OsOSCA1.2 had eleven TMs and the eighth hydrophobicity peak is composed of two TMs (**Fig. 1B**) based on hydropathy analysis and comparison of regions with the fungal homolog *Nectria haematococca* TMEM16 (NhTMEM16) (10). For convenience, we have kept the numbering convention of TMs consistent with NhTMEM16, and thus we refer to OSCA1.2’s additional N-terminal TM as TM0 (**Figs. 1C-D**). Despite a relatively low degree of sequence similarity, we later confirmed that OsOSCA1.2 shares significant structural homology to the TMEM proteins with respect to ten of the eleven transmembrane regions, corresponding to TMs1-10 in the mouse TMEM16A (mTMEM16A) structures.

TM0 threads from the extracellular N-terminal end of the protein through the membrane, linking to TM1 via a ~50 residue strand that is likely conformationally flexible. This portion of the protein is the only region not fully resolved in our density maps (**Figs. 1B-D**). A short helix on the cytoplasmic side then precedes TM1 and the C-terminal end of TM2 leads into the soluble cytosolic region of ~170 residues. The remaining helices represent the anoctamin domain, encapsulating the pore region for ion conductance. TMs3-4 are located on the outer edge of the transmembrane region and are tilted with respect to the membrane. TMs7-8 are shorter in length and are the only TMs that do not span the entire length of the membrane, with the connecting loop (residues 578-583) being embedded in the membrane and consisting of hydrophobic residues.

The soluble domain is located on the intracellular side of the channel joining TM2 and TM3 and makes important structural contacts with the C-terminus (**Figs. 1B-D**). A core globular domain comprises a four-stranded β-sheet buttressed by two short helices that interestingly forms a canonical RNA recognition motif (RRM) fold (12). Unlike true RNA binding RRM proteins, OsOSCA1.2 includes a fusion of a distinct 70-residue appendage between β-strands 2 and 3. These long, extended helical arms protrude out from the RRM domain and are located proximal to and in the plane of what would be the inner-leaflet side of the plasma membrane.

The dimer interface represents only a small percentage (~2.7%) of the surface area of each protomer, burying only ~1116 Å^2^ surface area comprised by interactions formed between the soluble domains (**Figs. 2A-B**) (13). Interface residues Q334, T335, Q336, Q337, T338, S339, L681, Q682, and E683 from both subunits likely make several hydrogen bonds and hydrophobic interactions (**Fig. 2C**). Interestingly, the TMs from each subunit do not cause significant interactions contributing to the dimer interface. The orientation and offset of the two halves of the dimer creates a large cavity between the two protomers, which, as predicted in other AtOSCA structures (6, 7, 14), is likely filled with lipids when embedded in the cell membrane.

**Figure 2.**
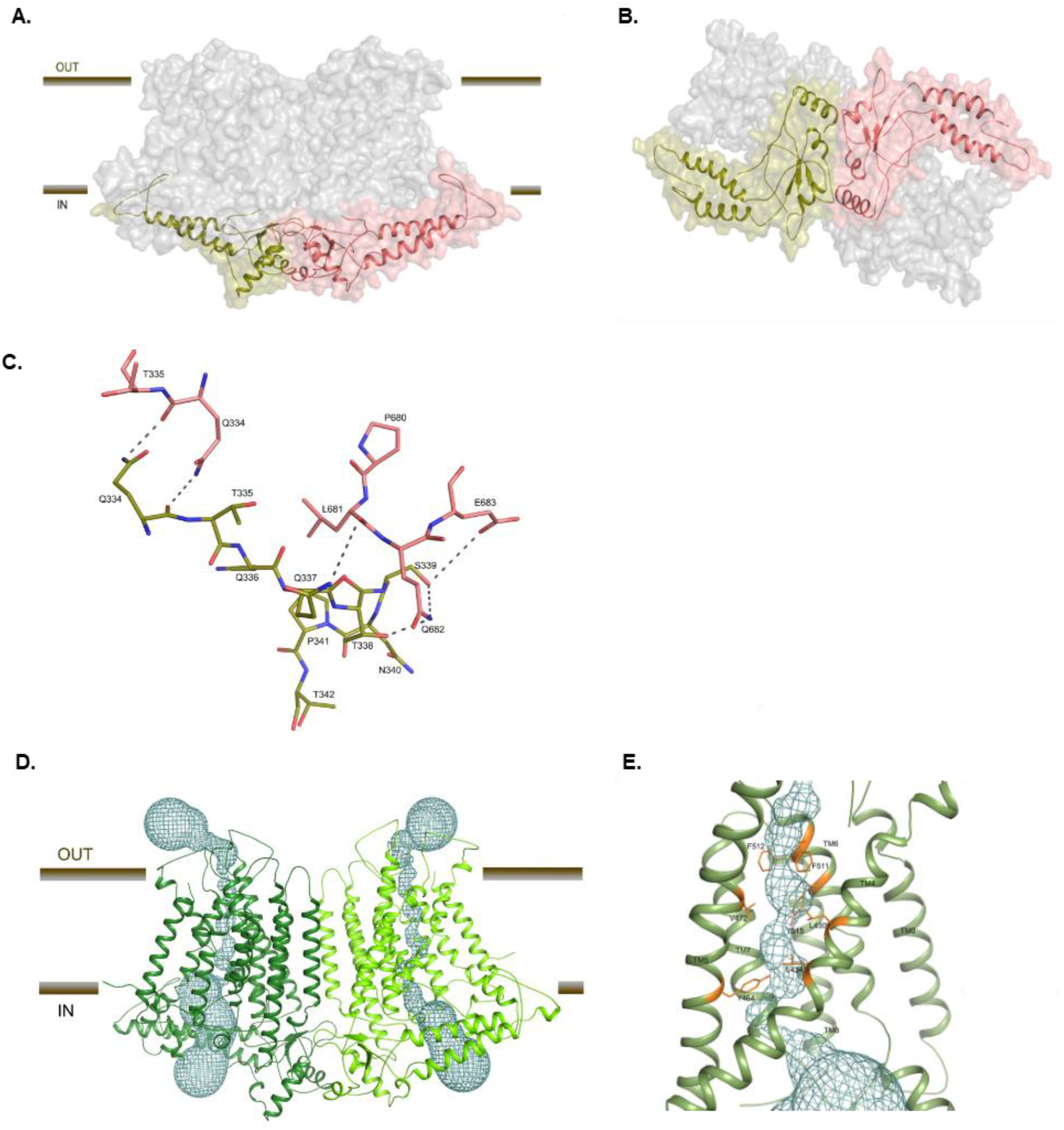
OsOSCA1.2 dimer interface and ion channel pore. (A) OsOSCA1.2 surface representation. The transmembrane domain is shown as gray and the cytoplasmic domain is colored red and yellow. (B) View of OsOSCA1.2 from the cytoplasmic side. (C) Dimer interface residues. (D) Location of the ion conductance pore in both subunits of OsOSCA1.2. The pore pathway is depicted in a cyan mesh. (E) Close-up view of the neck region, showing the residues ‘gating’ the pore.

The ion conductance pore of OsOSCA1.2 is contained within each monomeric subunit (**Fig. 2D**), and formed between TMs 3-7 as suggested by the topological similarities with mTMEM16A. Using the program HOLE to visualize the putative ion permeation pathway (15), the overall shape of the pore resembles an hourglass. The extracellular and an intracellular vestibule are bridged by a narrow neck region that is about 20 Å long through the membrane. The putative pore has an opening more than 12Å wide towards the extracellular side and narrows into the ‘neck’ region approximately 15 Å down the conduction pathway. The tightest juncture is ~0.8 Å wide, suggesting that this channel structure is in a closed conformation. The calculated pore profile predicts that the hydrophobic residues F511, F512, and Y515 on TM6 and V472 and Y464 on TM5 forms a gate that completely blocks the channel pore (**Fig. 2E**).

### OsOSCA1.2 computational dynamics

The DynOmics suite allows prediction and identification of candidate functional sites, signal transduction, and potentially allosteric communication mechanisms, leveraging rapidly growing structural proteomics data (16). The suite integrates two widely used elastic network models while taking account of the molecular environment like the lipid bilayer providing collective dynamics of structural resolved systems. We used DynOmics to do molecular dynamic simulations on our OsOSCA1.2 dimer model after embedding in the membrane, looking for regions that could potentially serve as functionally important sensors, broadcasters, and receivers (**Fig. 3A**). Our results revealed that the extended intracellular helical arms could communicate conformational perturbations, having the propensity to act as a broadcaster/receiver, extending to the central core sheet structure of the soluble domain and, more interestingly, TM6, which is proposed to be the ion gating helix in related structures (6, 7, 10, 11, 14).

**Figure 3.**
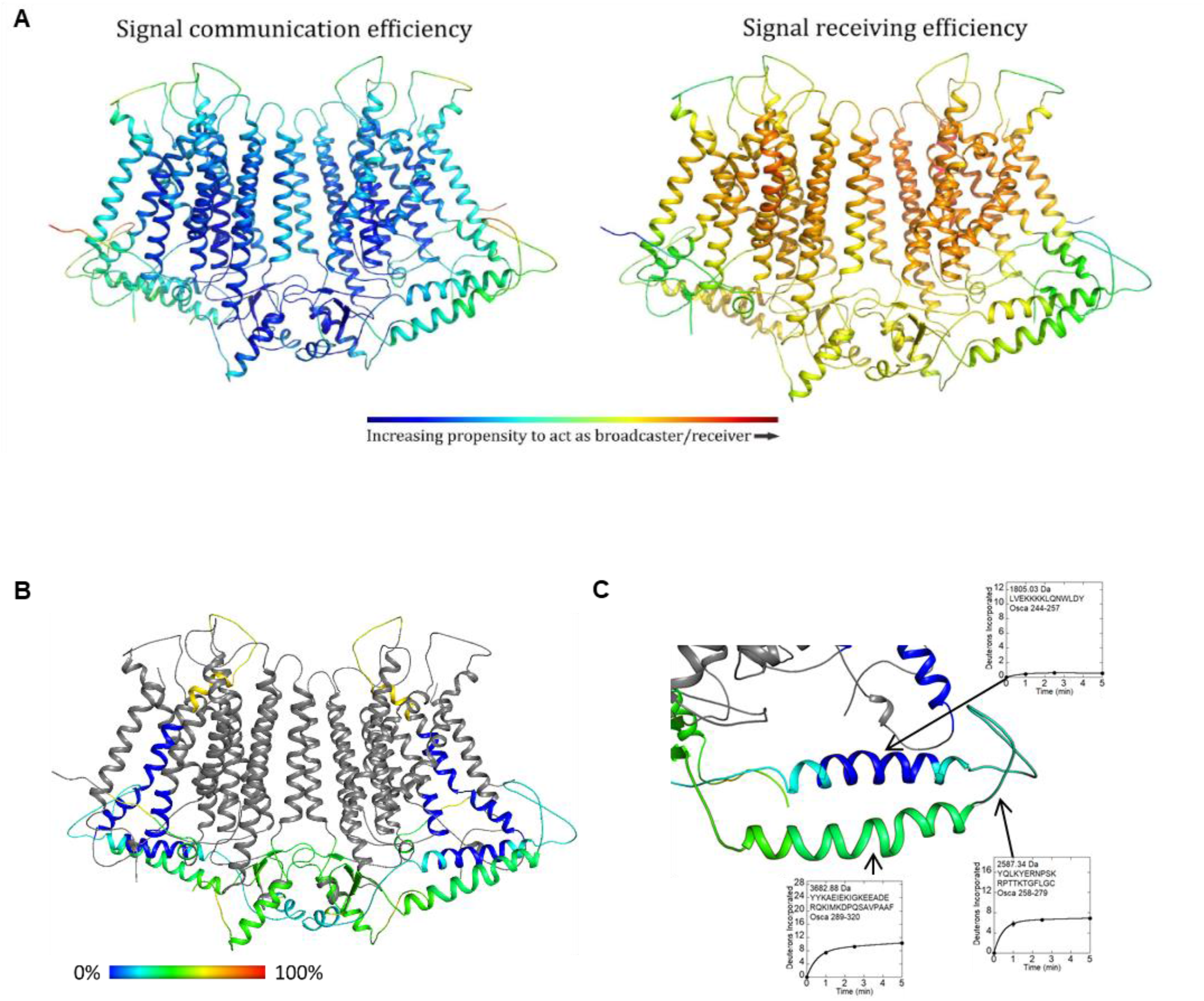
Computational and experimental dynamics of OsOSCA1.2. (A) Results of OsOSCA1.2 embedded in membrane using the Dynomics suite. Panels show a color-coded map superimposed on the model showing signal communication (left) and receiving (right) efficiency. Regions that are colored red are more active while those blue inactive with regards to molecular dynamics prediction. (B) Relative uptake after 5 minutes of exchange. Structure is color scaled and superimposed on model. Regions colored gray yielded no detectable peptide fragments (C) Close-up view of the extended and gating helix. Uptake plots for selected peptides are shown. Corresponding protein segments are outlined.

### OsOSCA1.2 experimental dynamics

In order to further understand and probe local conformational dynamics of OsOSCA1.2, we used HDXMS. This approach utilizes the exchange that occurs between protons linked to amide bonds and protons or deuterium nuclei from solvent molecules to provide experimental information regarding regional solvent accessibility and dynamics. When a protein is added to a solution containing excess D2O, hydrogen-deuterium exchange occurs most rapidly for protons which are exposed to solvent and unconstrained by intermolecular hydrogen bonds. Protease digestion and chromatographic separation facilitate the quantification of deuterium nuclei incorporated throughout the protein as measured by mass spectrometry (17). For the hydrogen-deuterium exchange to occur in well-folded regions, a protein must sample exchange-competent conformations which expose amide protons. Thus, the uptake of deuterium over time reflects the local dynamics that individual regions of a protein undergo in solution (18).

HDXMS measurements using detergent solubilized OsOSCA1.2 protein resulted in the identification of 32 peptides, which constitute 34.5% coverage of the molecule (**Figs. 3B** and S4A), including the helical arms that were predicted to dynamically couple to the presumed gating helix TM6 and most likely to be responsible for sensing lateral tension in the membrane. The helix closest to the inner leaflet side of the membrane was covered by two peptides (corresponding to residues 244-257 and 245-257). Deuterium incorporation profiles revealed that this region was tightly protected from exchange, indicative of rigid dynamics or association with a nearby surface (**Fig. 3C**). The following segment was also covered by two peptides (residues 258-279 and 258-286), which correspond to the C-terminal end of the protected helix and a nearby loop in our structure. This region was ~25% saturated with deuterium nuclei at the earliest measured time point of 1 minute, indicating rapid exchange associated with conformational flexibility. The remainder of this segment increased deuterium content by ~5% over 5 minutes suggesting conformational motions that gradually increased exposure to solvent. The helix farther from the membrane was covered by three peptides (residues 287-320, 289-320, 305-320) and similarly displayed rapidly-exchanging amides and ongoing deuterium exchange. Mass spectra from peptides corresponding to the unstructured loop and helix farther from the membrane all displayed bimodal deuterium uptake, which was more prominent among peptides corresponding to the loop (**Fig. S4B**). The ongoing dynamics in sharp contrast to the rigidity of the helix (residues 241-266) closer to the membrane. Despite being spatially and sequentially near each other, these two intracellular helices have very different dynamic properties.

### Topological insertion of OSCA in tA201 Cells

As the OSCA family inserts an addition helix, TM0, we investigated the orientation of the ion channel in the cell membrane in mammalian human embryonic kidney tsA201 cells. The topological prediction and structures suggested that the N- and C-termini of the molecule were on opposite sides of the membrane (**Fig. 1B**). We, therefore, made two C-terminal HA-tagged (YPYDVPDYA) cDNA constructs of OsOSCA1.2 and AtOSCA1 in a pcDNA3.1 vector. Human kidney cell-line tsA201 cells were transfected with both of these constructs (**Figs. S5A-B**) and, after 48 hours, the cells were stained with anti-HA antibody conjugated to Alexa488 without permeabilization (see Methods). Our results suggest that the C-terminus of the molecule was accessible only from outside the cell.

### Osmotic stress response of OsOSCA1.2

It was suggested that the orthologs of OsOSCA1.2 in *Arabidopsis*, AtOSCA1 and AtOSCA1.2, are osmotic stress responsive cation channels (1, 19). We, therefore, explored whether changes in the intracellular free Ca^2+^ concentration occur in response to osmolality changes in tA201 cells using a ratiometric fluorescent indicator, Fura-2AM (**Fig. S6**). No obvious differences were observed between control and OsOSCA1.2-expressing tA201 cells in hypoosmotic (168 mosmol kg^-1^) and hyperosmotic (627 mosmol kg^-1^) calcium responses (**Figs. S6A-B**, n > 23 OsOSCA1.2-expressing cells). Rapidly induced fluorescence ratio changes were observed in response to exogenous ATP in the same OsOSCA1.2-expressing tA201 cells as controls (**Fig. S6A**), suggesting that Fura-2 can efficiently report intracellular calcium changes in these experiments.

## Discussion

OsOSCA1.2 shares overall protein fold and topology with other recently determined homologous structures from *A. thaliana* (6, 7, 14). A superposition of these structures with OsOSCA1.2 showed a significant difference (rmsd ~3-4Å) for the pore-lining helices (TM3-7) along with TM0 and TM8 (**Fig. 4A**). When comparing intracellular soluble domains, the extended helical arms of OsOSCA1.2 had noticeable differences compared to that of AtOSCA1 (**Fig. 4A**). These differences are likely due to a combination of conformational flexibility inherent in the detergent solubilized protein and structural difference between species.

**Figure 4.**
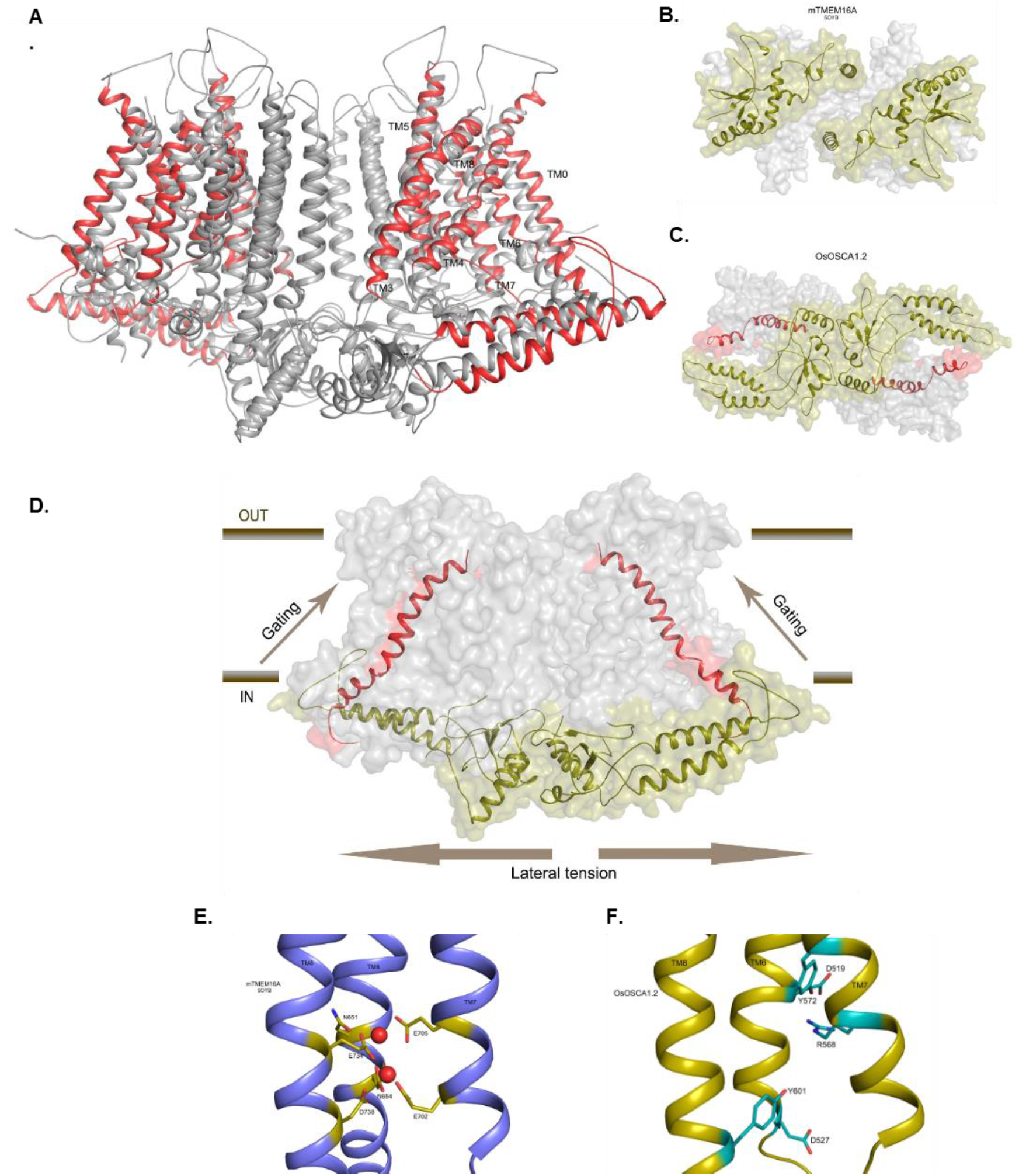
Structural comparisons of OsOSCA1.2 with other TMEM and OSCA structures. (A) Superposition of OsOSCA1.2 and AtOSCA1 (PDB: PYD1). TMs 1, 2, 9, and 10 (shown in gray) close to lipid-filled cleft are nearly superimposable and have little relative movement. Pore-lining helices (TMs 3-7) showed significant movement along with TM0 and TM8 (shown in red). (B) The mTMEM16A soluble domains from intercellular side are separated. (C) OsOSCA1.2 intracellular soluble domains are together and communicate with channel gating helix TM6. (D) General mechanism of OsOSCA1.2 shown in the plane of the lipid membrane. Lateral tension on the inner leaflet side of the lipid bilayer causes a conformational change in the extended helices of the soluble domain, which is coupled to the gating helix TM6 opening pore. (E) Calcium binding site residues of mTMEM16A. Calcium ions are shown as red spheres. (F) The corresponding region of OsOSCA1.2 with charged and polar residues are shown in cyan.

OsOSCA1.2 also shares structural homology to the TMEM family (mTMEM16A, NhTMEM16) when comparing monomeric transmembrane domain regions. However, they differ significantly in the regions of extra- and intra-cellular loops and domains as well as the intermolecular packing arrangement of the respective dimers. The dimer interface of mTMEM16A buries less surface area (~2%) compared to OsOSCA1.2 and most of the interactions are mediated through the TM domains. The intracellular domains of mTMEM16A and nhTMEM16, which are formed by the N and C-termini of the molecule, do not contribute to the formation of the dimer (**Fig. 4B**). In contrast, OsOSCA1.2 dimerizes mostly through interactions formed between the opposing intracellular soluble domains. This distinct dimeric packing resulted in a more pronounced offset between protomers, that is ~20 Å wider for OsOSCA1.2 compared to NhTMEM16 or mTMEM16A. In our OsOSCA1.2 structure, the extended helical arm in the intracellular soluble domain make hydrophobic contacts with the loop connecting the gating helix TM6 (**Fig. 4C**), suggesting a possible role for the helical ‘arms’ in sensing the membrane tension and, in turn, transmitting these conformational/mechanical changes to gate ion conductance (**Fig. 4D**). This feature is, of course, missing in TMEM16 channels as Ca^2+^ ions control gating.

Our OsOSCA1.2 structure likely represents the non-conducting state conformation as no tension, pressure or osmolality mismatch was applied between the intracellular and extracellular sides of the OsOSCA1.2 protein during cryo-EM sample preparation. Indeed, the tightest juncture in the pore is ~0.8 Å wide (**Fig. 2D-E**). Our DIBs study suggests that the reconstituted OsOSCA1.2 protein could conduct ions (**Fig. S1E**). Interestingly, when the OsOSCA1.2 structure was compared to mTMEM16A, we found a π-helical turn at TM6 near the ‘neck’ region of the pore that might be associated with ion gating and channel opening in a similar fashion (Ca^2+^ activated π-to-α transition) to that observed in mTMEM16A (11, 20). The regulatory Ca^2+^ binding site composed of acidic and polar residues (E702 and E705 from TM7, E734, and D738 from TM8, and N651 and E654 from TM6 in mTMEM16A) are well conserved in the TMEM16 family (**Fig. 4E**). However, when compared to the corresponding same region in OsOSCA1.2, the negatively charged residues D519 and E527 on TM6 and polar residue R568 on TM7 locate spatially different (**Fig. 4F**). Therefore, OsOSCA1.2 will likely not specifically bind calcium ions in this region.

Computational dynamic studies using our OsOSCA1.2 model and experiments using hydrogen-deuterium exchange with detergent solubilized OsOSCA1.2 protein provided a molecular structural basis of how OsOSCA1.2 couples osmotic stress to induce ion channel gating in the membrane spanning region. Taken together, both studies predict and suggest that the extended helical arms (residues 241-266) have the mechanical rigidity and propensity to act as a broadcaster/receiver, transmitting conformational changes caused by lateral tension in the membrane to TM6 (**Figs. 3, 4F**), which is important for gating ion conductance. In addition, information from HDXMS revealed the presence of bimodal deuterium exchange throughout the OsOSCA1.2 (**Figs 3B-C and S4B**), most prominently within the helical arms (residues 258-320) and some extracellular loops (residues 489-511). Bimodal exchange is indicative of multiple correlated unfolding processes occurring in the observed regions (21). Interestingly, in each peptide where bimodal peaks were observed, the two peaks remained equal in intensity over the entire course of the experiment, suggesting at least two distinct conformational states occupied by the molecular ensemble at equilibrium in the resting state.

Several electrophysiological studies have used mammalian cells over-expressing OSCA channels to measure conductance gated by direct mechano-transduction or pressure (1, 4, 6). Interestingly, we found that the C-terminus and presumably the entire cytosolic domain (residues 191-363) were on the outside when OSCA channels were over-expressed in mammalian cells (**Fig. S5**). This finding suggests and could explain why channel opening due to changes in ion concentration may be impaired. As previous studies suggested Ca^2+^ conductance, we further probed ion flux using FURA2 using these cells as proxy for calcium but found no changes relative to controls for either hyper-or hypo-osmotic conditions (**Fig. S6**). It remains unclear whether or not these ion channels can be gated by changes in ion concentration. Interestingly, analysis of the taxonomic distribution of different OSCA fragments suggests that TM0 is restricted to plants and that the cytosolic domain (residues 191-363; cytoL2) is probably distributed similarly to the rest of OSCA family (**Fig. S7**-9 and **Table S1**). In fungi and plants, the N-terminus of TM0 is predicted to be on the outside with the osmo-sensing cytosolic domain inside the cell (**Fig. 1B**). OSCA may be inserted in the membrane differently between mammalian and plant/fungi cells and may be an important consideration in their functional study.

Several important questions remain regarding these OSCA channels and their role in crops. For example, the specificity of endogenous ions conducted by OsOSCA1.2 or by other members within their greater family is still unknown. It will certainly be a challenge to assign function to all of these proteins individually as these channels are members of large gene families. Although we present a structure of OsOSCA1.2 along with computational and experimental dynamics, the detailed function mechanism(s) coupling lateral tension in the membrane by OsOSCA1.2 to channel gating remains to be addressed in future studies. These studies will certainly be challenging given the transient nature of channel gating.

## Acknowledgments

We thank Valorie D. Bowman for assistance during data collection at the Purdue Cryo-EM facility supported by NIGMS grant U24 GM116789. This work was funded by NSF PGRP IOS-1444435 and in part supported by the National Institute of Health (GM060396).

## Methods

### Expression and Purification of OsOSCA1.2

We cloned OsOSCA1.2 (GenBank Kj920372.1) and made TEV protease cleavable green fluorescent protein (GFP) fusions into the pPICZc vector, and tested expression in *Pichia pastoris*. Expression vectors were linearized using *PmeI* and electroporated into competent *P. pastoris* KM71H cells (Life Technology). The resulting transformants were cultured and induced in small scale to screen for target expression based on the intrinsic GFP fluorescence of cells and also from an anti-His western blot of whole cell lysate. OsOSCA1.2 was found to show both high levels of expression, and desirable properties during purification (described below) and was therefore chosen for further characterisation. Yeast clones selected for their high expression of OsOSCA1.2 were grown in minimal glycerol (4%) media, supplemented with 0.4% phosphoric acid and 0.024% trace metals at 28°C in a New Brunswick BioFlo 415 (Eppendorf). The pH of the media was titrated to pH 5 prior to inoculation and adjusted during the vegetative growth phase using 50% ammonium hydroxide. The dissolved oxygen (DO) was maintained at 10% minimally through-cascaded agitation until a DO spike occurred. The fermentation culture was then induced at pH 5 by slow methanol addition for 16-18 hours.

Cells were harvested and resuspended in cold lysis buffer (20 mM Tris-HCl pH 8.0, 100 mM NaCl, 15% glycerol, 23.4 mM leupeptin, 7 mM E-64, 4 mM chymostatin, 14.5 mM pepstatin A, 1 mM PMSF, 25 mM benzamidine), and they were lysed by a single passage through a cell disruptor (TS-Series, Constant Systems, Inc.) at 40,000 psi. Cellular debris were removed by centrifugation (12,500 x g, 20 minutes, 4°C) and the supernatant continued onto a 38,400 x *g* spin for 4 hrs to fractionate the plasma membrane. The membrane fraction was resuspended in lysis buffer and frozen at −80°C.

Membranes were solubilized with 1% n-dodecyl-β-D-maltopyranoside (DDM) and 0.1% sodium cholate for ~90 min at 4°C. Insoluble material was removed by centrifugation (38,400 x g, 4 °C for 60 minutes) and 15 mM imidazole was added to the supernatant before batch binding to Ni-NTA agarose resin (Qiagen). The bounded resin was sequentially applied to a gravity column housing and washed with buffer A (20 mM HEPES pH 8.0, 150 mM NaCl, and 0.03% DDM, 0.003% cholesteryl hemisuccinate) and an imidazole gradient was applied. Bound target protein was eluted with buffer A containing 300 mM imidazole, concentrated to ~8 mL, desalted (HiPrep 26/10, GE Healthcare) and subjected to TEV protease digestion for 12 hours at 4°C. TEV digested sample was reapplied to Ni agarose (Qiagen) to rebind the TEV protease and the C-terminal His-GFP tag. The collected OsOSCA1.2 was then concentrated to ~1 mL and ultraspun at 95,000 rpm (TLA120.1 rotor) for 15 minutes at 4°C. The sample was then applied to a Superdex 200 increase size-exclusion column (GE Healthcare) pre-equilibrated with 20 mM HEPES pH 8.0, 150 mM NaCl, 0.06% n-undecyl-β-D-maltopyranoside 0.2 mM tris(2-carboxyethyl)phosphine, and 0.01% cholesteryl hemisuccinate, and run at 4°C. Peak fractions off the size-exclusion chromatography column were checked using sodium dodecyl sulfate polyacrylamide gel electrophoresis (SDS-PAGE) and directly snap frozen at a concentration of ~3 mg/mL.

### Determination of the molecular mass of OsOSCA1.2 using SEC-MALLS

Size-exclusion chromatography coupled to multi-angle laser light scattering (SEC-MALLS) was performed using a Superdex 200 Increase 10/300 GL size exclusion column (GE Life Sciences) connected in series to a miniDAWN TREOS light scattering detector and an Optilab T-rEX refractive index detector (Wyatt Technology, Santa Barbara, CA, USA). Purified OsOSCA1.2 was injected onto the column with 20 mM HEPES pH 8.0, 150 mM NaCI, 0.03% n-dodecyl-β-D-maltopyranoside, 0.2 mM TCEP,0.001% sodium cholate and 0.02% cholesteryl hemisuccinate and run at 0.4 mL/min. The elution was monitored in-line with three detectors, and the molecular weights of the protein-micelle complex, the micelle and the protein were calculated using ASTRA v.6 software (Wyatt Technology) in conjugate mode as previously described (22).

### Functional reconstitution of OsOSCA 1.2 into droplet interface bilayers (DIBs)

Lipids of *E. Coli* extract polar (Avanti #100600) were dried under nitrogen and vacuum desiccated for one hour, before resuspending in reconstitution buffer (10 mM HEPES, pH 7.4, 150 mM KCl) to a final concentration of 10 mg/ml. The resuspended lipids were incubated for a minimum of 20 min followed by addition of 0.1% DM. The mixture was allowed to sit at room temperature for 30 min followed by bath sonication for 5 cycles of 1 minute sonication and 2 min on ice. Purified OsOSCA1.2 was then mixed at a protein-lipid ratio of 1:500 (w/w) with the detergent saturated liposomes. The protein was reconstituted by removal of detergents by the detergent-dilution method (23). OsOSCA1.2 containing proteoliposomes were resuspended to a final lipid concentration of ~5mg/ml, extruded through a 100 nm filter, and stored at −80°C until use. Ion channel reconstitution into droplet interface bilayers (DIBs) was done as detailed elsewhere (24–26). Briefly, a lipid asymmetric droplet-droplet (~200 nl) configuration (27) was obtained by placing OsOSCA1.2 containing proteoliposomes in *E. coli* extract polar lipids on the agar coated head-stage Ag electrode and POPC: POPS: DOPA (1:1:0.5; Avanti #850457, #840034, #840875) liposomes on the reference electrode. The droplets were incubated in a hexadecane (Sigma # H6703) medium for 5-10 min to allow formation of monolayers on each droplet. The two droplets were then brought into contact with each other and the formation of bilayers was monitored using a triangular wave protocol. Incorporation of ion channels into the bilayer were detected as discrete fluctuations in current amplitude under voltage clamp conditions. Data were acquired using a Dagan 3900A amplifier and pCLAMP 10 software (Molecular Devices, Sunnyvale, CA). Data were filtered at 1 kHz and sampled at 250 kHz and using a Digidata 1440A. Downward deflections traces represent inward currents *(cis* to *trans)*, whilst upward deflections represent outward currents *(trans* to *cis)*. Single channel data analysis was performed using Clampfit 10 (Molecular Devices). The cation ionic concentrations of all solutions were verified experimentally via conductively coupled plasma emission spectrometry and their ionic activities were calculated using GEOCHEM-EZ (28). All experiments were performed at room temperature.

### EM Data Collection

Quantifoil 1.2/1.3 Au(Quantifoil Micro Tools GmbH, Germany) or C-Flat 1.2/1.3 300 (Protochips, Raleigh, NC) mesh grids were glow discharged for 30 seconds at 110mA (Emitech). Four microliters of OSCA1.2 at a concentration of 1.8 mgs/ml was applied to the grids, blotted for 2.5s at a relative humidity of 100% and plunge frozen in liquid ethane using a FEI Vitrobot Mark 2 (FEI Company, Hillsboro, OR). Two image sets were collected. The first data set was collected using defocus phase contrast on an FEI Tecnai F30 microscope(FEI Company, Hillsboro, OR) operating at 300kV with a K2 Summit camera(Gatan, Inc., Pleasanton, CA) at a nominal magnification 31,000x in super resolution mode with a pixel size of 0.636Å using SerialEM software(Mastronarde Group, Boulder, CO). A total of 40 frames at 200 ms per frame were recorded for each image at a camera dose rate of 8 electrons/pixel/s. A total of 342,910 particles covering a defocus range from −0.8 to −2.8 microns were used to determine an initial 6.0 Å resolution map that was utilized to build a poly alanine model (**Table 1**). A higher resolution data set was collected using a FEI Titan Krios equipped with a Volta Phase Plate, GatanEnergy Filter and a K2 summit camera (Gatan, Inc., Pleasanton, CA). Data were collected at a nominal magnification of 105,000x in super resolution mode and a total of 64,096 individual particle images at a fixed target defocus of −0.5 micron defocus were used to determine the structure at 4.9 Å resolution (**Table 1 and Fig. S2**).

**Table 1.** Cryo-EM data collection, 3D reconstruction and model building.

| **Data Collection and Processing** |  |  |
| --- | --- | --- |
| Microscope | FEI Tecnai F30 | FEI Titan Krios |
| Voltage (kV) | 300 | 300 |
| Camera (mode) | Gatan K2 Summit<br>(40 frame super-res movies) | Gatan K2 Summit<br>(40 frame super-res movies) |
| Target Defocus ($\mu\text{m}$) | -0.8 to -2.8 | -0.5<br>Volta Phase Plate |
| Pixel size ($\text{\AA}$) | 0.6355 | 0.6920 |
| Imposed symmetry | C2 | C2 |
| Electron dose ($\text{e-}/\text{\AA}^2$) | 45.5 | 40.8 |
| Initial particle images | 499,157 | 169,655 |
| Final particle images | 342,910 | 64,096 |
| Map resolution ($\text{\AA}$) FSC 0.143<br>Max local resolution ($\text{\AA}$) | 6.0 $\text{\AA}$<br>4.8 $\text{\AA}$ | 4.9 $\text{\AA}$<br>4.5 $\text{\AA}$ |
| **Model Building and Refinement** |  |  |
| Map sharpening B factor ($\text{\AA}^2$) | -800 | -530 |
| Protein residues (expected) | 1388 (1424)* | 1388 (1424) |
| R.M.S. Z score<br>Bond Lengths (# Z>2)<br>Bond Angles (# Z>2) |  | 0.30 (2)<br>0.44 (6) |
| Validation<br>MolProbity score<br>Clashscore<br>EMRinger score<br>Poor rotamers (%) |  | 1.98<br>7.99<br>0.89<br>0.00 |
| Ramachandran plot<br>Favored (%)<br>Outliers (%) |  | 90.1<br>0.0 |

### EM Data Processing

The Tecnai F30 data set consisted of a total of 9,691 micrographs in 3 groups were selected for initial processing after motion correction using MotionCor2 and CTF estimation with Gctf. Non dose-weighted micrographs were used for CTF estimation, and dose-weighted micrographs for all other processing. Approximately 1,000 manually picked particles from each group were used to generate 2X binned templates (2.542 Á pixel size) which were used for autopicking in Relion. Micrographs. Autopicked particles were manually screened, and 499,167 particles extracted for further processing in cryoSPARC. 2D classification and selection yielded 342,910 particles which were then used for initial model construction and auto-refinement. Auto-refinement and masking with C2 symmetry yielded a map with 6.0 Á resolution by GSFSC corrected for the effects of masking. Local resolution estimation in cryoSPARC indicated that the core regions have resolutions ranging from 4.5 Á to 6.0 Á.

The Titan Krios data set consisted of a total of 2,408 micrographs that were motion corrected using MotionCor2 and CTF’s estimated using Gctf. Results were imported into Relion 2.1, A total of 1,126 corrected micrographs were selected for further processing after screening for excessive motion, and poor or poorly estimated CTF’s. 1,134 particles were manually picked, classified in 2D, and the selected templates used to auto-pick 372,278 particles. Further screening resulted in selection of 650 micrographs containing 169,655 particles for additional processing. 2X binned (2.76 Á pixel size) particles were extracted and processed through 2 rounds of 2D classification and selection, resulting in 64,096 remaining particles. 3D autorefinement with C2 symmetry using these particles and an initial model from the previous defocus contrast refinement yielded a model with 7.4 Á resolution. Reextraction with unbinned pixels and subsequent refinement led to no improvement in resolution at this stage. 3D classification into 10 classes and subset selection yielded 5 subsets (best single class, best 2 classes, best 5 classes, and all classes) which were used for another round of re-extraction and auto-refinement. The best resolution resulted from using all 64,096 particles and was unchanged at 7.4 Á. Masking and post-processing resulted in an estimated resolution of 6 A. At this point, the 64,096 extracted particles were transferred to cryoSPARC, and subsequent processing performed in cryoSPARC. 3D auto-refinement with C2 symmetry using all 64,096 particles and an initial model constructed using a subset of 16,438 selected particles resulted in an GSFSC estimated resolution of 4.9 Á. The auto-refined, unsharpened map was further sharpened with a B-factors ranging from −350 to −600 out to a cutoff of 3.5 Á for modelling and a map with a B-factor of −530 was used for subsequent model building and refinement.

### Model building and Refinement

An initial polyalanine model was built using the 6.0 Å resolution map with multiple rounds of real space refinement in Phenix/COOT (29, 30). In order to determine the absolute hand at this resolution, the initial and inverted model were used for molecular replacement using X-ray diffraction data set that extended to 9 Å in resolution. Only one model provided a solution to the MR search. Subsequent use of this initial model and the observation of the helical hand in the 4.9 Å resolution map further confirmed the correctness of the assigned hand. The full atomic model was built into the higher resolution map using multiple rounds of building and real-space refinement in COOT and Phenix. The density maps within the transmembrane region were of sufficient quality to readily identify large aromatic side chains (**Fig. S3**) and helped to confirm the correct sequence registration. Comparison to the recently determined structures of the AtOSCA1.2 (6, 14) further confirmed the correctness of our model despite the lower calculated overall resolution of our map.

### Image Processing

Motion-corrected projections with pixel size 1.271 Å (F30) and 1.384 Å (Titan Krios), with and without dose-weighting, were constructed using MotionCor2 (31) with 2X binning and grouping. The CTF estimation was performed using Gctf (32) followed by manual selection to remove micrographs with poor or incorrectly fit CTF, poor astigmatism and contamination. Manual and semi-automated particle picking was done using RELION 2.1 (33), followed by sorting and another round of manual over-reading to remove low quality micrographs. Subsequent refinements were carried out in RELION or cryoSPARC (34). Local resolution estimation was performed using cryoSPARC or ResMap (35).

### Hydrogen-deuterium exchange mass spectrometry (HDXMS)

HDXMS measurements were made using a Synapt G2Si system (Waters Corporation). Deuterium exchange reactions were carried out by a Leap HDX PAL autosampler (Leap Technologies, Carrboro, NC). Deuterated buffer was prepared by lyophilizing 10 mL of 20 mM HEPES, pH 8.0, and 150 mM NaCl. Lyophilized buffer was resuspended in 10 mL of 99.96% D2O immediately before use, to which was added powdered n-undecyl-β-D-maltopyranoside to a final concentration of 0.06% and cholesterol hemisuccinate to a final concentration of 0.01 %. Each deuterium exchange time point (0 min, 1 min, 2.5 min, 5 min) was measured in triplicate. For each measurement, 4 μL of protein at a concentration of 5 μM was mixed with 36 μL of D2O buffer at 25 °C. Deuterium exchange was quenched by combining 35 μL of the deuterated sample with 65 μL of 0.1% formic acid and 3M guanidinum-HCl for 1 min at 1 °C. The quenched sample was then injected in a 50 μL sample loop and digested by an inline pepsin column (Pierce, Inc.) at 15 °C. The resulting peptides were captured on a BEH C4 Vanguard precolumn at a flow rate of 400 μL/sec, separated by analytical chromatography (Acquity UPLC BEH C4, 1.7 μM, 1.0 × 50 mm, Waters Corporation) using 7-85% acetonitrile in 0.1% formic acid over 7.5 min, and analyzed in a Waters Synapt G2Si quadrupole time-of-flight mass spectrometer following electrospray injection.

Data were collected in Mobility, ESI+ mode, mass acquisition range of 200-2000 (m/z), scan time 0.4 s. Continuous lock mass correction was performed using infusion of leu-enkephalin (m/z = 556.277) every 30 seconds (mass accuracy of 1 ppm for calibration standard). For peptide identification, data were collected in MS^E^ (mobility ESI+) mode. Peptide masses were identified following triplicate analysis of 10 μM OsOSCA1.2, and the data were analyzed using PLGS 2.5 (Waters Corporation). Peptide masses were identified using a minimal number of 250 ion counts for low energy peptides and 50 ion counts for their fragment ions. The following parameters were used to filter peptide sequence matches: minimum products per amino acid of 0.2, minimum score of 7, maximum MH+ error of 5 ppm, and a retention time RSD of 5%, and the peptides had to be present in two of the three ID runs collected. After identification in PLGS, peptides were analyzed in DynamX 3.0 (Waters Corporation). Deuterium uptake for each peptide was calculated by comparing the centroids of the mass envelopes of the deuterated samples with the undeuterated controls. To account for back-exchange and systematic autosampler sample handling differences, the uptake values measured at the 1 min time point were divided by 0.79. The longer 2.5 min and 5 min deuteration time point deuteration values were divided by 0.75. Data were plotted as number of deuterons incorporated vs time. The Y-axis limit for each plot reflects the total number of amides within the peptide that can possibly exchange. Each plot includes the peptide MH+ value, sequence, and sequential residue numbering.

### Production of OsOSCA1.2/AtOSCA1 stable expression cell-line

An epitope HA tag (YPYDVPDYA) was introduced onto the 5’ end (HA-OsOSCA1.2) or the 3’ end (OsOSCA1.2-HA, AtOSCA1-HA) of the full-length OsOSCA1.2 or AtOSCA1 cDNAs by PCR. The cDNA was amplified using the *PfuUltra II* Fusion HS DNA Polymerase (Agilent Technologies, Inc., Santa Clara, CA). PCR products were then subcloned into the pcDNA3.1 vector (Thermo Fisher Scientific, Waltham, MA). The cDNA inserts were verified by sequencing (GENEWIZ, South Plainfield, NJ). After linearization of the vectors with *Pvu I* enzyme, the vectors were transfected to tsA201 cells (ECACC) using Lipofectamine LTX with Plus Reagent (Thermo Fisher Scientific, Waltham, MA). Stable cell-lines were selected 48 h post-transfection with 1mg/ml of Geneticin (Thermo Fisher Scientific, Waltham, MA).

### Immunofluorescence assay

OsOSCA1.2/AtOSCA1 expressing cells were plated on poly-L-lysine-coated glass coverslips in 24-well plates. After 48 hours, cells were fixed with 4% paraformaldehyde/PBS. And then cells were blocked with 3% BSA/PBS for 30 min. Expression of OsOSCA1.2/AtSCA1 were detected with Alexa Fluor 488 anti-HA (16B12) antibody (BioLegend, San Diego, CA) in PBS, 1% BSA/PBS for 1 h. Samples were visualized on a fluorescence microscope (EVOS Cell Imaging Systems, Thermo Fisher Scientific).

### [Ca^2+^]_i_ Imaging in tA201Cells

Intracellular Ca^2+^ concentration changes in tA201 cells in were observed using the ratiometric Ca^2+^ indicator dye, Fura-2 (36). The mammalian cells were transfected with the *pCDNA3.1* (control) or the *pCDNA3.1-OsOSCA1.2-HA* vector were cultured on poly-L-lysine coated glass bottom 35 mm dish for 16 to 24 hrs. Cells were loaded with 5 μM Fura-2AM (F1221, Invitrogen, Eugene, OR) in loading buffer (~286 mosmol kg^-1^; 130 mM NaCl, 3 mM KCl, 0.6 mM MgCl_2_, 0.1 mM CaCl_2_, 10 mM glucose, 10 mM HEPES, adjusted to pH 7.4 with NaOH) and kept in the dark for 45 min, washed twice by assay buffer (~286 mosmol kg^-1^; 130 mM NaCl, 3 mM KCl, 0.6 mM MgCl_2_, 2 mM CaCl_2_, 10 mM glucose, 10 mM HEPES, and adjusted pH to 7.4 by NaOH) for 5 min each time, and then incubated with 1 mL assay buffer for Ca^2+^ imaging. For hypoosmotic treatments, 1 mL hypertonic buffer (3 mM KCl, 0.6 mM MgCl_2_, 2 mM CaCl_2_, 10 mM glucose, 10 mM HEPES, and adjusted to pH 7.4 using NaOH) was added into the original 1 mL assay buffer resulting in a final osmolality ~168 mosmol kg^-1^ of the incubation buffer. For ATP treatments, 20 μl of 1 mM ATP was added into the 2 mL ~168 mosmol kg^-1^ incubation buffer resulting a final ATP concentration of 10 μM. For hyperosmotic treatments, 1 mL of hypertonic buffer (650 mM sorbitol, 130 mM NaCl, 3 mM KCl, 0.6 mM MgCl_2_, 2 mM CaCl_2_, 10 mM glucose, 10 mM HEPES, and adjusted to pH 7.4 with NaOH) was added resulting in a final osmolarity ~627 mosmol kg^-1^ in the incubation buffer. The osmotic concentrations of the buffers were determined by using a Wescor 5500 Vapor Pressure Osmometer. Time-resolved Fura-2 imaging was performed using an Eclipse TE300 inverted microscope equipped with a Plan Fluor 40x/1.30 Oil objective DIC H °°/0.17 WD 0.2 (Nikon, Tokyo, Japan), a ET-Fura2 filter set 79001 (EX340x, EX380x, ET510/80m; Chroma, Bellows Falls, VT), a Mac 2002 System automatic controller, a Cool SNAP HQ camera (Photometrics, Tucson, AZ), and guided by the MetaFluor software version 7.0r3 (Molecular Devices, Sunnyvale, CA). Fluorescence images excited at 340 nm or 380 nm were collected at 200 ms exposure every 5 seconds. Emission ratios for 340/380 nm excitations in cells were processed and analyzed using Fiji (37).

## Data availability

Cryo-EM maps of OsOSCA1.2 has been deposited to the Electron Microscopy Data Bank under accession codes XXXX and XXXX. Atomic coordinates of OsOSCA1.2 have been deposited in the PDB under ID XXXX. All other data are available upon request to the corresponding author(s).

**Figure S1.**
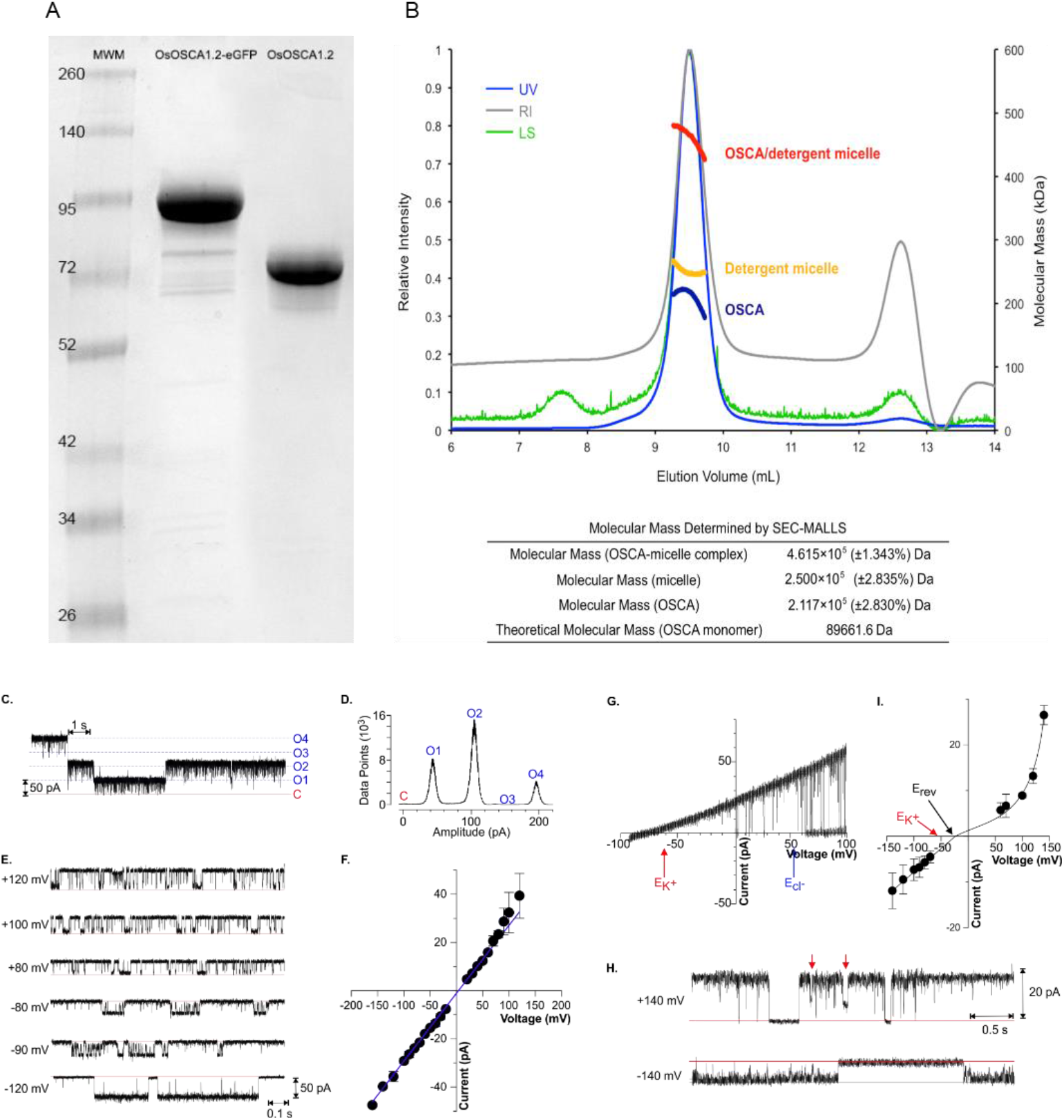
Purification and Reconstitution of OsOSCA1.2. (A) SDS-PAGE (sodium dodecyl sulfate polyacrylamide gel electrophoresis) of purified OsOSCA1.2 protein with and without GFP tag. (B) Purified βDDM solubilized OsOSCA1.2 was analysed using size exclusion chromatography in combination with multi-angle laser light scattering (SEC-MALLS). Chromatographs show ultraviolet (UV, blue), refractive index (RI, gray) and light scattering (LS, green) detector readings normalized to the peak maxima (left axis). The thick lines indicate the calculated molecular masses (right axis) of the complete protein/detergent complex (red), as well as the contributions of the detergent (yellow) and protein components (blue) throughout the elution peaks. SEC-MALLS analysis suggests that purified OsOSCA1.2 exists as a dimer in solution. The molecular mass values of the OSCA-micelle complex, the micelle and the protein as determined by SEC-MALLS. The molecular weight of OsOSCA1.2 was calculated from the amino acid sequence. (C-H) Reconstitution of OsOSCA1.2 proteoliposomes into droplet interface bilayers resulted in discrete single channel currents. (C) Example traces of four OsOSCA1.2 channels incorporated into a lipid bilayer recorded in symmetric 150 mM KCl conditions at a holding potential of +140 mV. Upward deflections indicate channel opening representing outward currents (*trans* to *cis*). The zero current (closed state) and conducting (open state) levels are indicated by the red solid and blue dotted lines, respectively. Time and current scales are shown on the top and bottom left corners. (D) Relative histograms illustrating the close and open states distributions for the full-length recordings illustrated on the left. The zero current level (closed state) and conducting (open) state labelled on top correspond to those levels shown on traces on the left. (E) Single channel recordings from a bilayer containing a single active OsOSCA1.2 channel. Recordings were obtained in symmetrical conditions (150:150mM KCl) in response to the holding potential indicated on the left of each trace. The zero current levels are indicated by the red solid lines. At positive potentials, upward deflections indicate channel opening representing outward currents (*trans* to *cis*); at negative potentials downward deflections indicate channel opening representing inward currents (*cis* to *trans*). Time and current scales for all traces are shown on the right bottom margin of the last trace. Note the different time scale relative to the traces shown in (C). (F) Single channel current amplitude as a function of voltage obtained from recordings as for those shown in F. In symmetrical 150 mM KCl conditions the channel showed no sign of current rectification, with a unitary conductance of 284 + 2 pS (n=3 bilayers), as determined from a linear regression for values between +120 and – 160 mV. (G) current-voltage ramp (−100 to +100 mV / 1.5 s) of an OsOSCA1.2 channel in asymmetrical 15:150 mM KCL (pH 7.4) *cis:trans* conditions. Four consecutive ramps are superimposed. The values for the theoretical Nernst potential for K^+^ (E_K_+) and Cl^-^ (E_C^-^_) are indicated by the arrows. (H) Example of single channel recordings from a bilayer containing a single active OsOSCA1.2 channel in asymmetrical conditions (15:150mM KCl) in response to the holding potential indicated on the left of each trace. The zero current levels are indicated by the red solid lines. The red arrows on top of the +140 mV illustrates a brief closure of about 50% the full current amplitude. This 50% state was rarely resolved as long-lasting events as those illustrated by the arrows, but rather appear as a fast flickery behaviour (also see traces in part E). (I) Single channel current amplitude as a function of voltage for OsOSCA1.2 under asymmetrical 15:150 mM KCL conditions. The current to voltage relationship was built from steady-state recordings of single-channel activity as those exemplified in part H. The unitary conductance of the inward current and reversal potential (E_rev_) were determined by fitting of a linear regression of the inward (i.e. negative) currents. E_rev_ (black arrow) was −26 mV and the slope (conductance) 103 + 4 pS. The theoretical Nernst potential for K^+^ (E_K_+) was −54 mV and is indicated by the red arrow.

**Figure S2.**
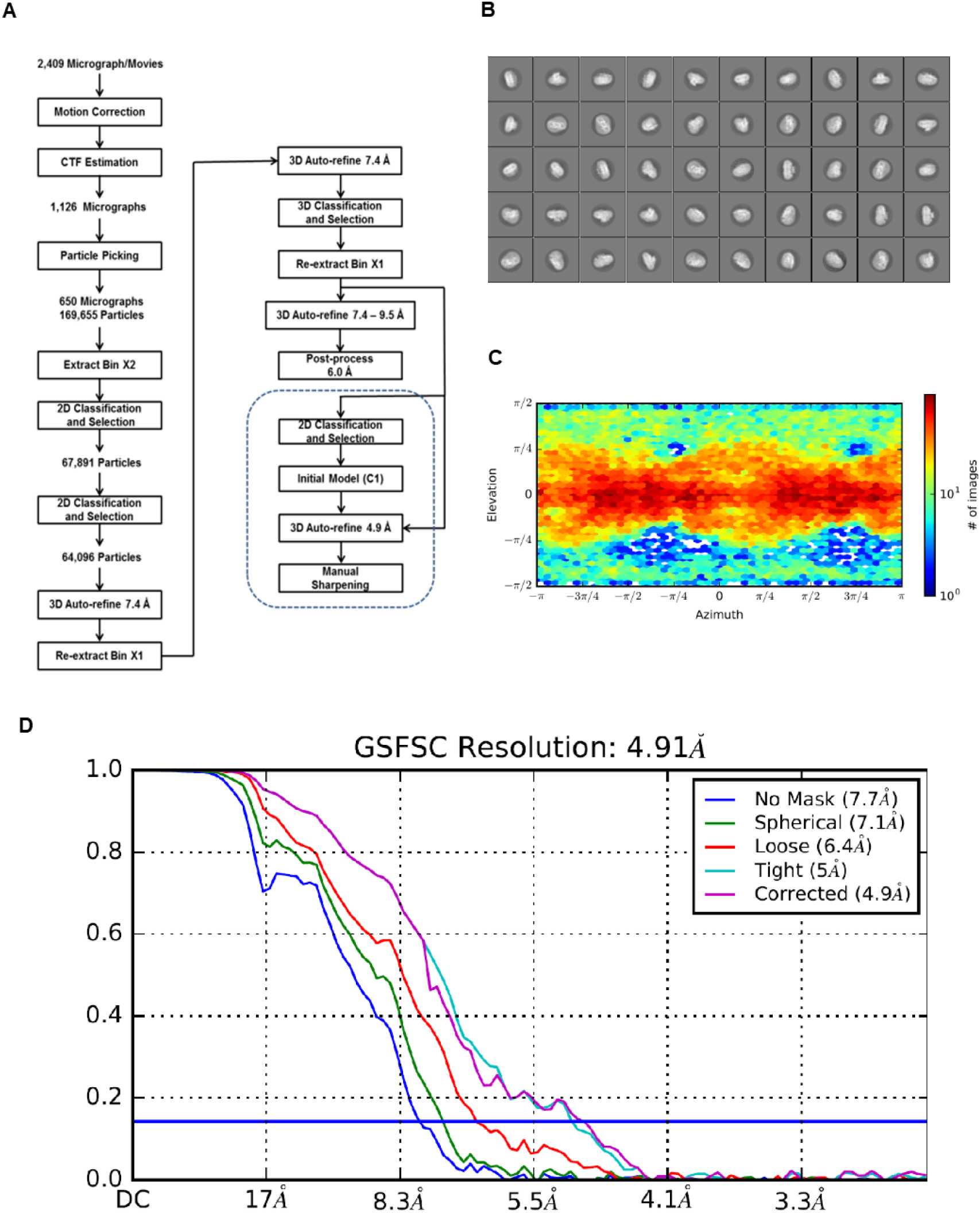
Cryo-EM of OscaA1.2. (A) Data processing flow chart for the 4.9 Á VPP map. (B) 2D-class averages. (C) Angular distribution of particles contributing to the final 4.9 Á map. (D) GSFSC plots of unmasked and masked maps.

**Figure S3.**
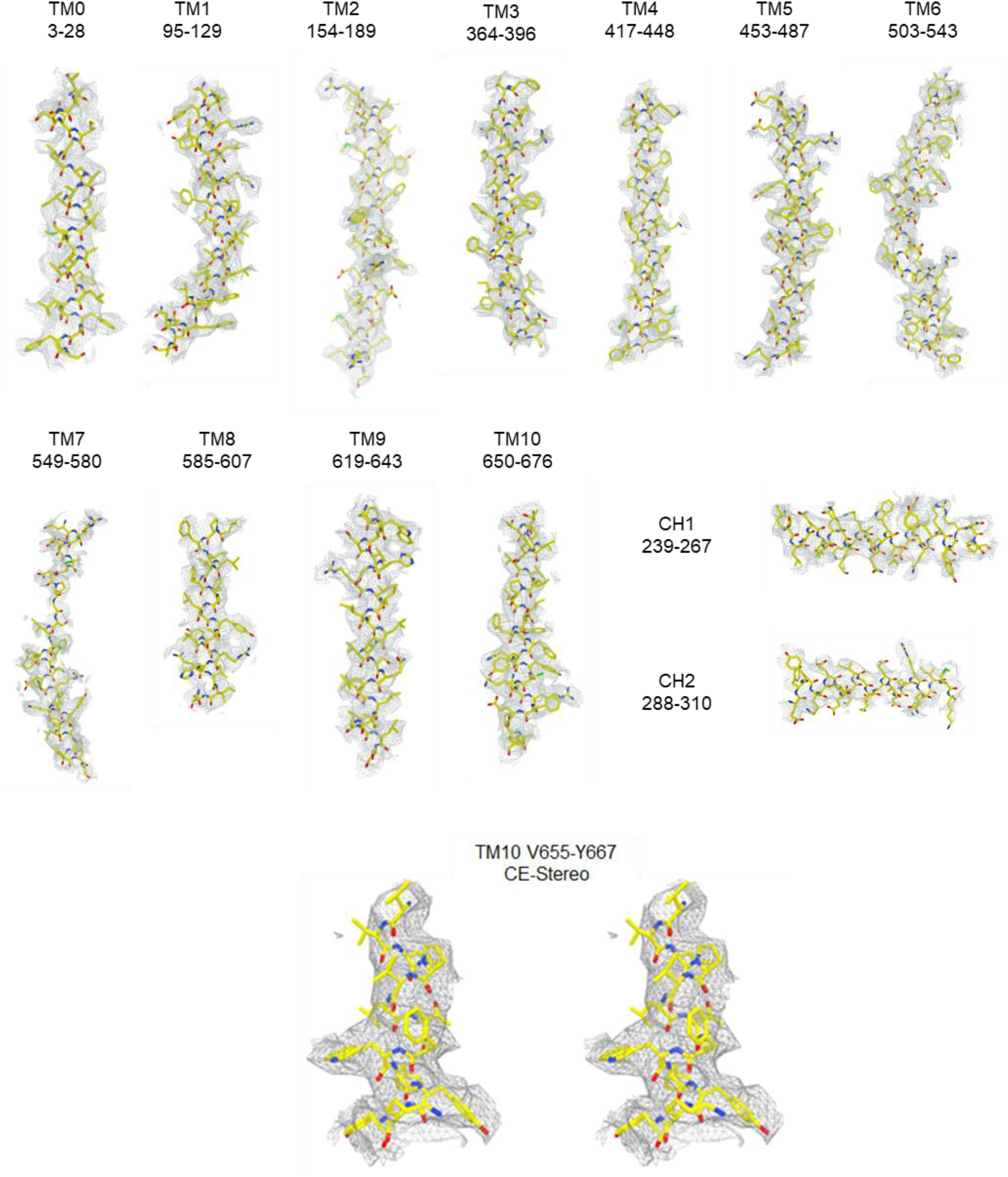
Representative density of OscA1.2. Shown are density maps for all 11 TM helices (TM0-TM10) and the cytoplasmic helices (CH1-CH2) of the soluble domain. A cross-eyed stereo-view of TM10 demonstrating the quality fit of large aromatic sides chins used to sequence registration.

**Figure S4.**
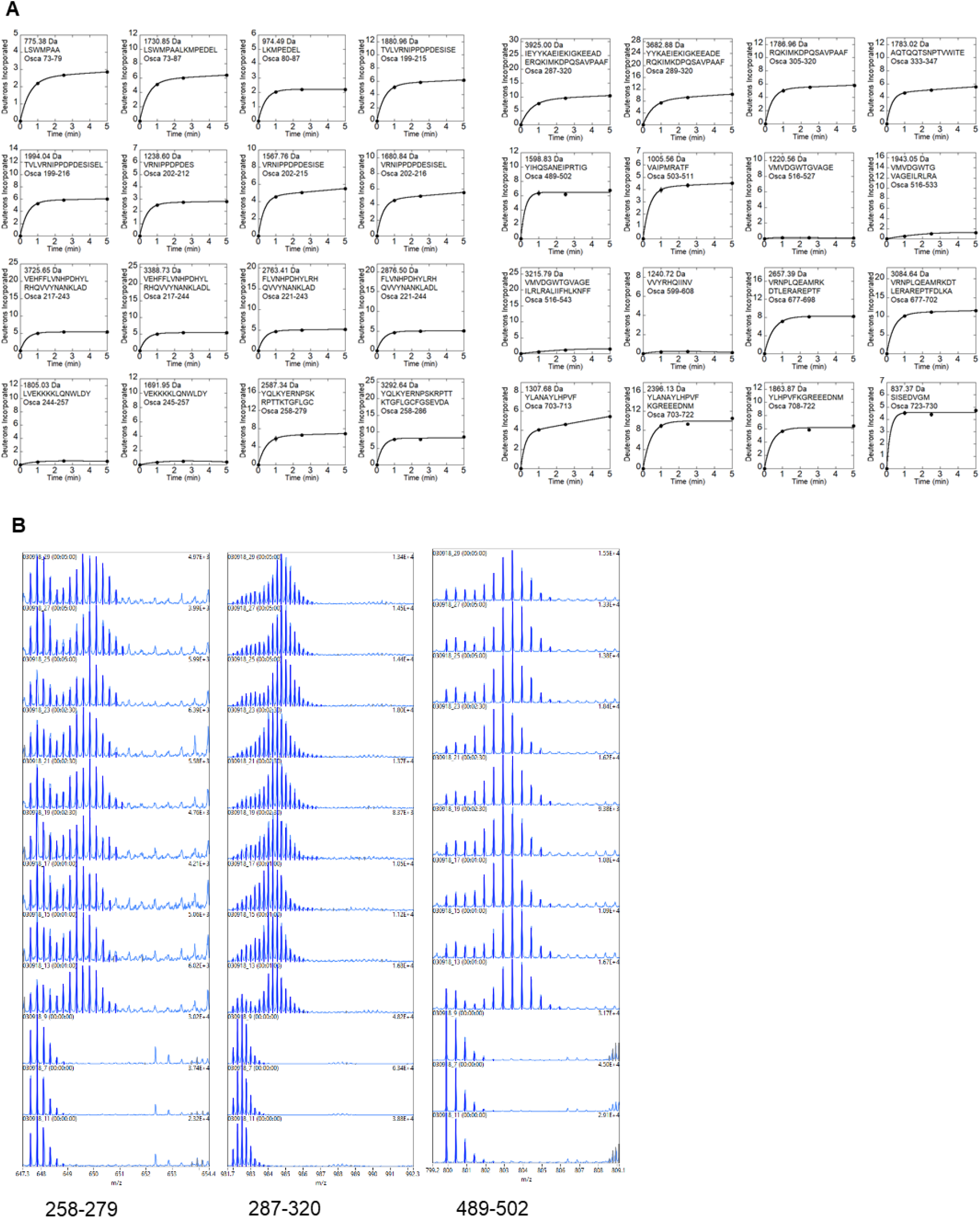
HDXMS Deuterium uptake plots. (A) Deuterium uptake plots for all peptides identified following HDXMS. (B) Representative mass spectra displaying bimodal deuterium uptake.

**Figure S5.**
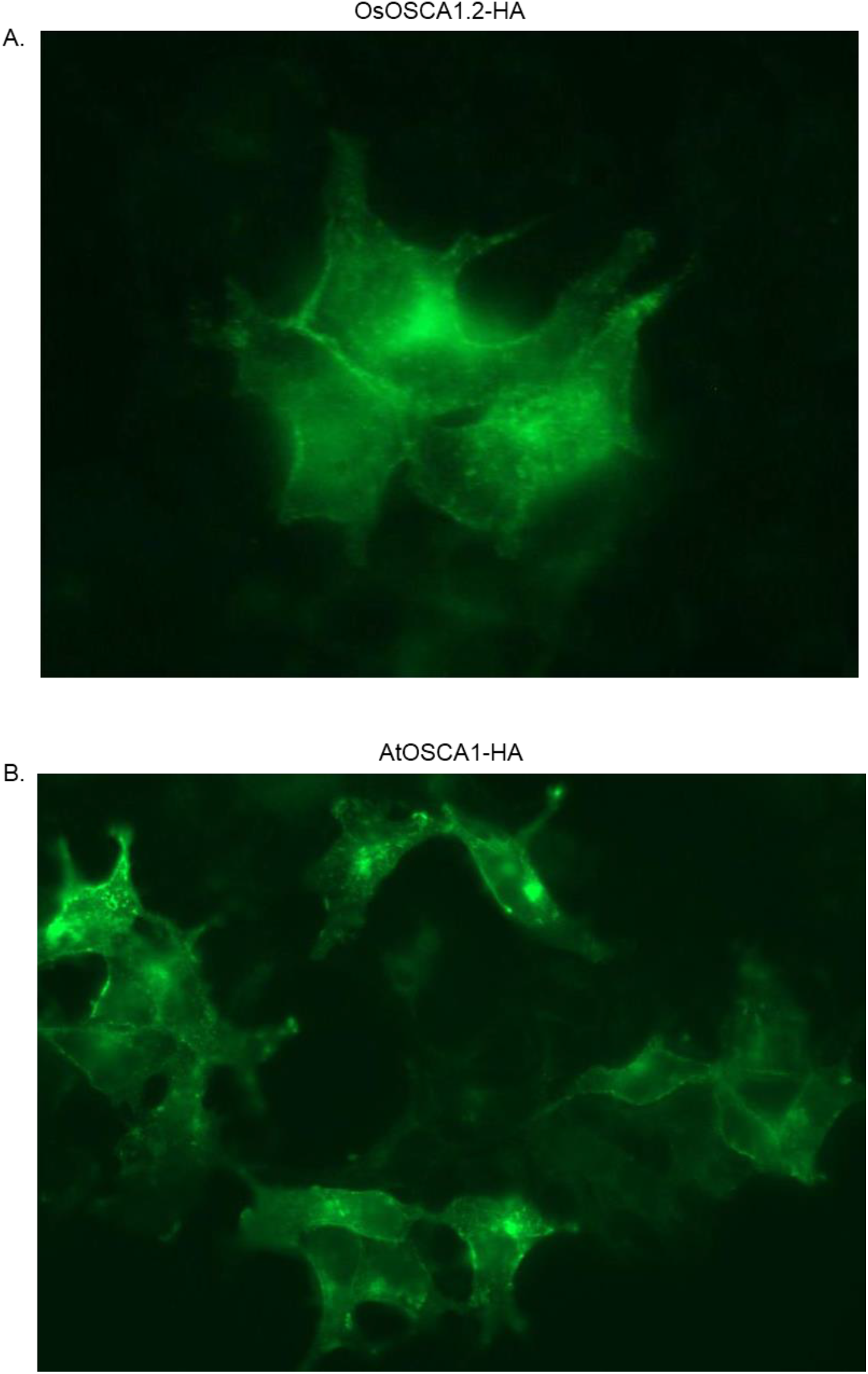
Expression of OsOSCA1.2/AtOSCA1 with C-terminal HA-tag in mammalian tsA21 cells. Immunostain of anti-HA antibody with Alexa 488 suggests exposure of C-terminal HA-tag on the outside of (A) OsOSCA1.2-HA and (B) AtOSCA1-HA. Cells were not permeabilized.

**Figure S6.**
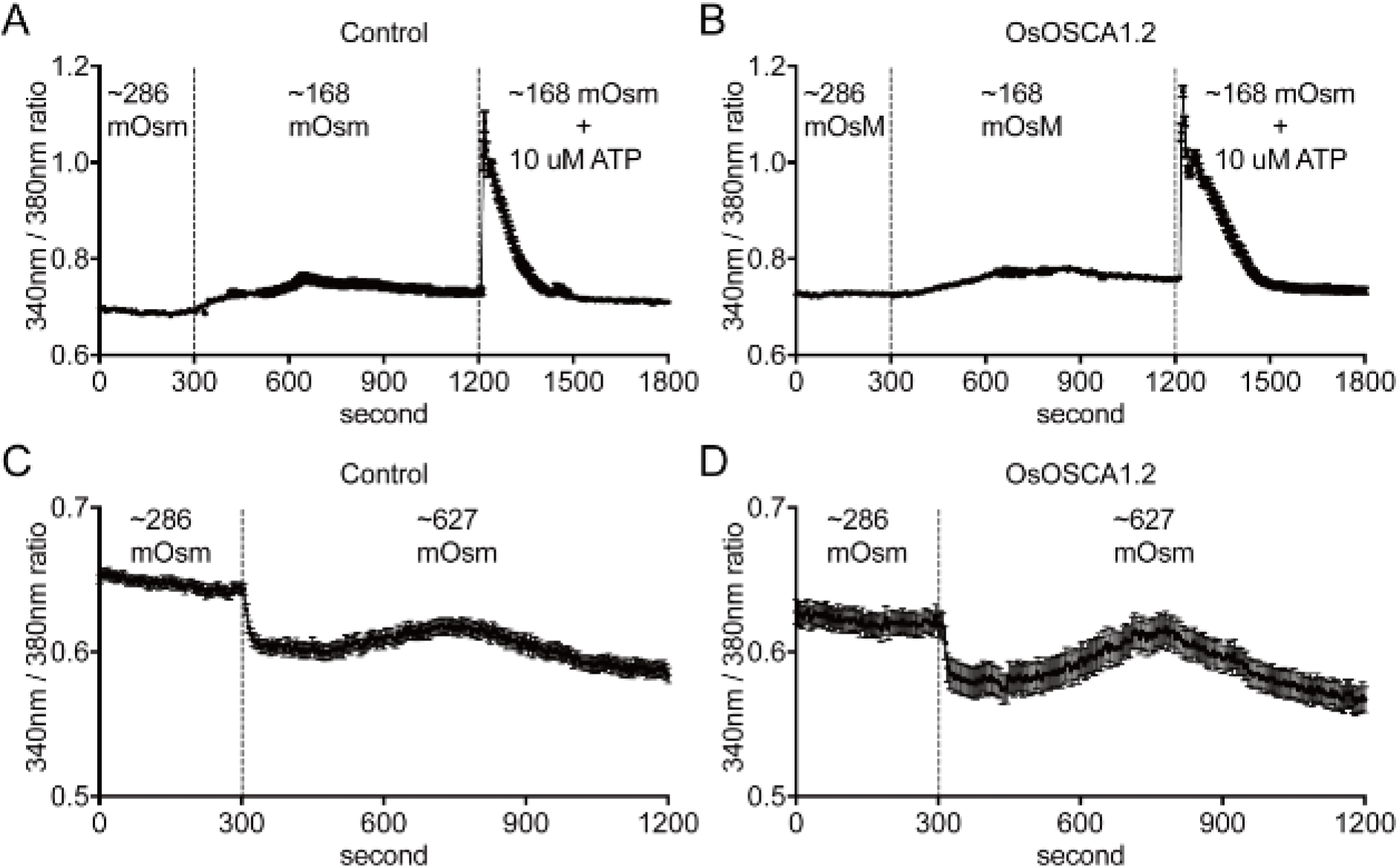
OsOSCA1.2 does not alter calcium responses in tsA21 cells under osmotic stresses. Time-resolved intracellular calcium concentration changes were analysed in tsA21 cells transformed with the pCDNA3.1 vector (control) or OsOSCA1.2 as indicated in each panel using a ratiometric fluorescent calcium indicator, Fura-2AM. The ratios of fluorescence intensities in response to excitation at 340 nm and 380 nm were calculated in individual cells and average traces of cells are shown (A and B). tsA21 cells were exposed to hypo-osmotic stress by shifting the osmolality of the bath solution from ~286 mOsm to ~168 mOsm and followed by adding 10 μM ATP. (C and D) tsA21 cells were exposed to hyper-osmotic stress by shifting the osmolarity of the bath solution from ~286 mOsm to ~627 mOsm. Data represent mean ± SE. n = 45 cells in A, 49 cells in B, 39 cells in C, and 23 cells in D.

**Figure S7.**
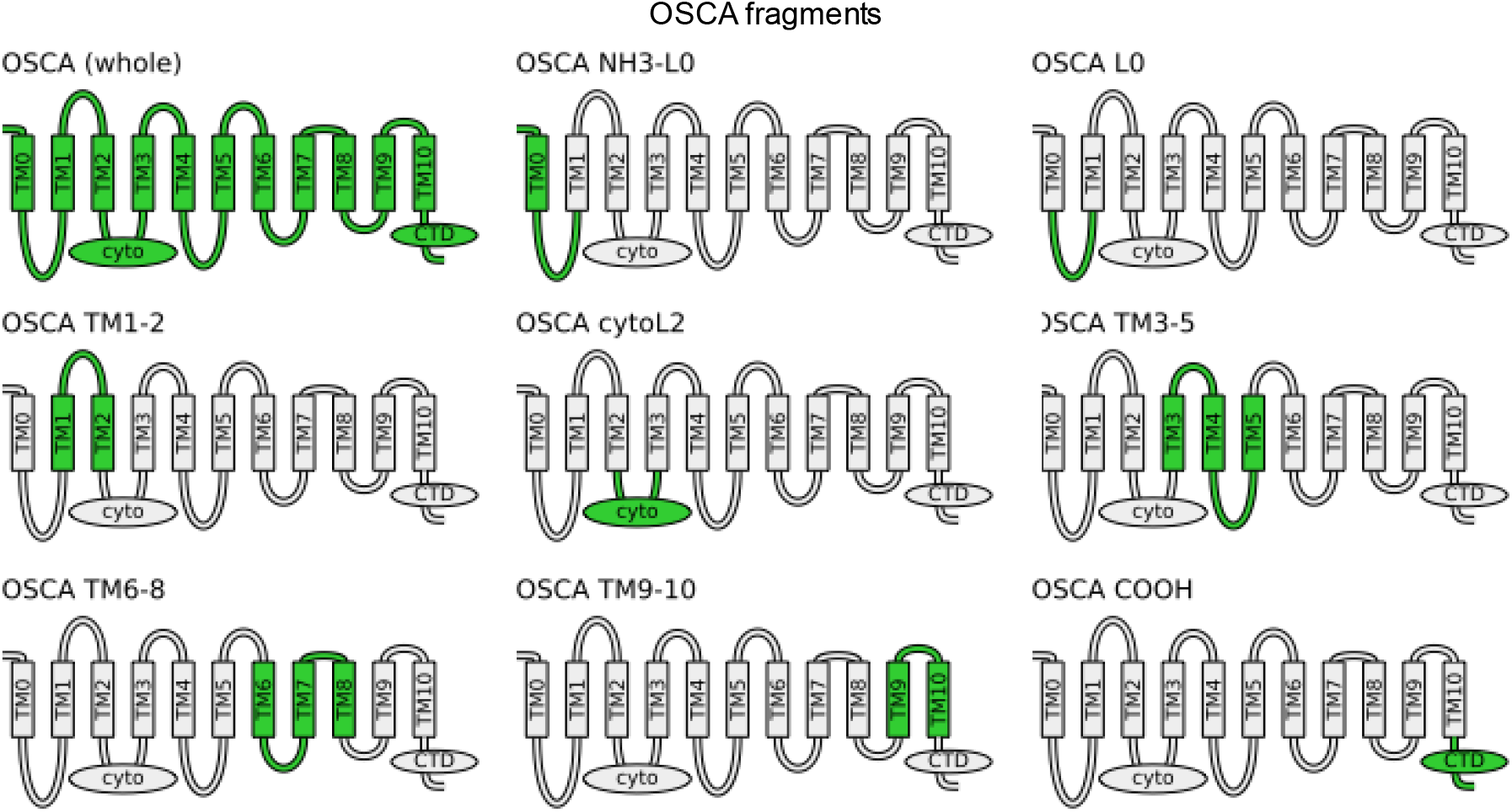
OSCA Fragments Selected for Homolog Searches. OSCA fragments selected for BLAST searches are shown in green while the remainder of the protein is shaded light gray. Due to its extremely short length (23aa), fragment NH3-TM0 was discarded in favor of NH3-L0, which includes loop 0. Two fragments containing L0 (NH3-L0 and L0) were selected to probe for the possible origin of TM0.

**Figure S8.**
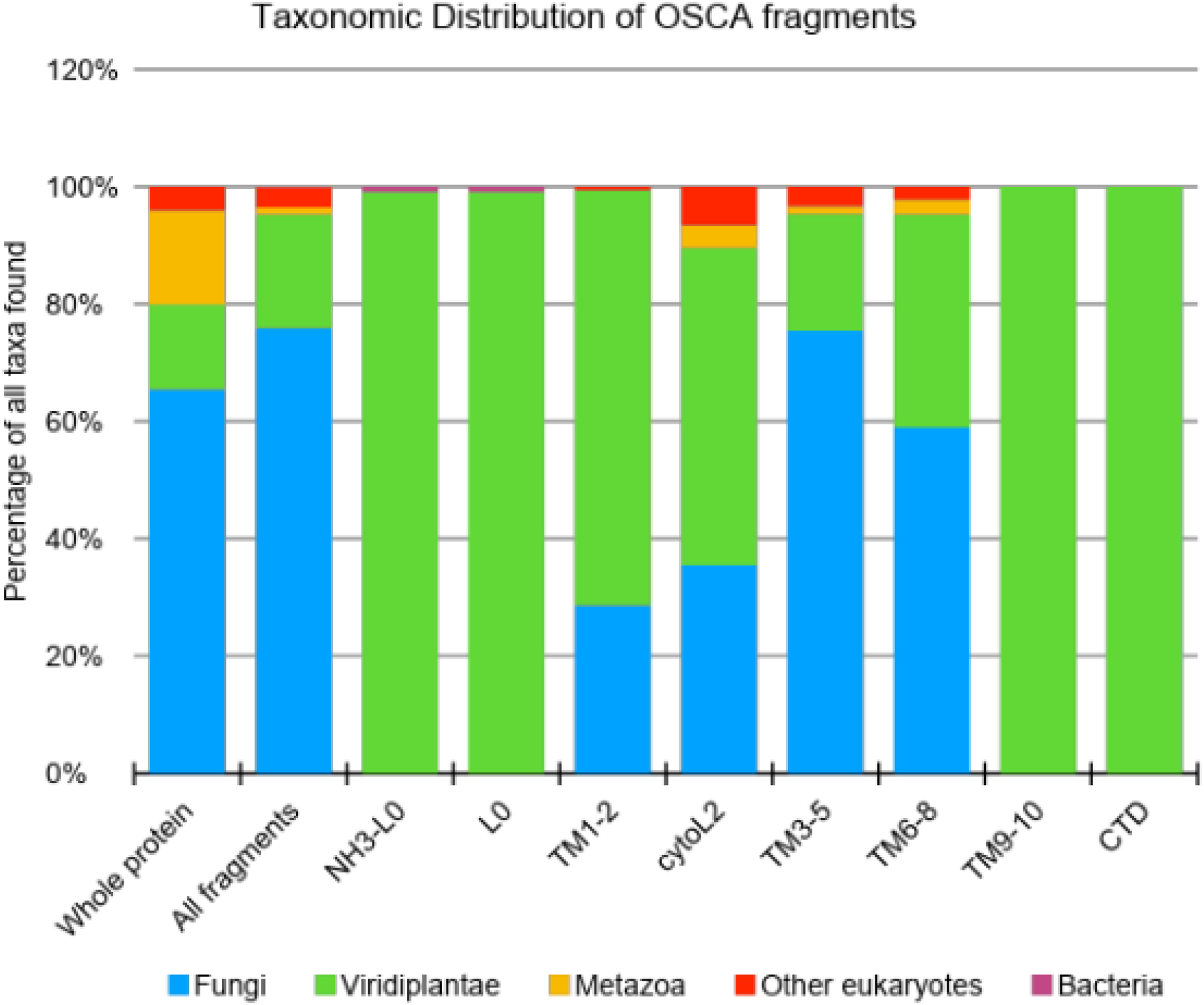
Taxonomic distribution of OSCA fragments. Columns 3 to 10 correspond to different OSCA fragments and show the relative frequency of the taxonomic groups identified in **Table S1**. Plants appear to be the only organisms with sequences similar to TM0, TM9, TM10, Loop 0, and the disordered C-terminal domain.

**Figure S9.**
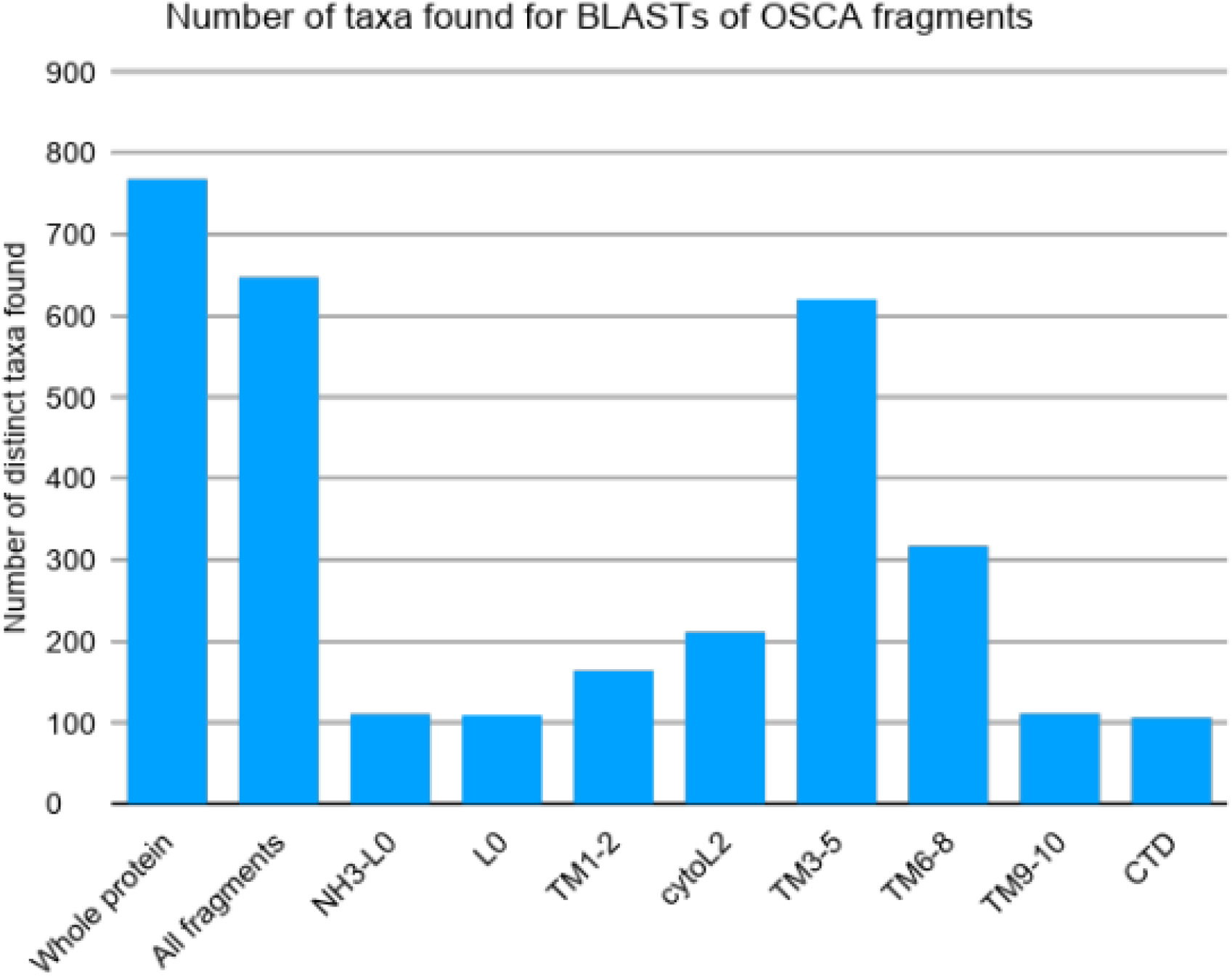
Raw frequency of taxa groups containing the different OSCA fragments. These are the counts used to generate the relative frequencies presented in Table S1 and Figure S9. Note that NH3-L0 and L0 hit the hypothetical enterococcal protein PWS22870 despite matching fewer taxa.

**Table S1:** Taxonomic distribution of selected OSCA Fragments. To narrow down the possible regions associated with OSCA’s osmo/mechanosensing behavior, the primary sequence of OsOSCA1.2 was split into eight fragments (**Figure S8**). Each fragment was BLASTed against the NCBI’s non-redundant protein database (E-value < 10^−5^ and query coverage ≥ 80%) and the taxonomic information associated with each hit was extracted. Each column corresponds to a separate fragment. The column labelled “All fragments” consists of the union of taxa after merging all the data. Interestingly, one small hypothetical protein (PWS22870) from *Enterococcus faecium* was found with high significance (E-value < 10^−40^; identity > 70%) and high coverage (88%) for the fragments containing TM0 and L0. However, as PWS22870 does not hit any other bacterial proteins, it cannot be ruled out that this is potentially the result of a sequencing error or a contaminating DNA fragment.

| Taxonomic Distribution of selected osOSCA fragments |  |  |  |  |  |  |  |  |  |  |  |
| --- | --- | --- | --- | --- | --- | --- | --- | --- | --- | --- | --- |
|  |  | Whole | All fragments | NH3-L0 | L0 | TM1-2 | cytoL2 | TM3-5 | TM6-8 | TM9-10 | CTD |
| BLAST stats |  |  |  |  |  |  |  |  |  |  |  |
|  | Query length | 766 | N/A | 92 | 69 | 96 | 172 | 118 | 103 | 53 | 94 |
|  | Hits | 3702 | 5017 | 1025 | 1025 | 1861 | 2606 | 3968 | 2978 | 1123 | 888 |
| Taxonomic stats (%) |  |  |  |  |  |  |  |  |  |  |  |
|  | Eukaryota | 100.0% | 99.8% | 99.1% | 99.1% | 100.0% | 100.0% | 100.0% | 100.0% | 100.0% | 100.0% |
|  | Fungi | 65.5% | 75.9% | 0.0% | 0.0% | 28.7% | 35.5% | 75.5% | 59.0% | 0.0% | 0.0% |
|  | Viridiplantae | 14.5% | 19.3% | 99.1% | 99.1% | 70.7% | 54.0% | 19.8% | 36.3% | 100.0% | 100.0% |
|  | Metazoa | 16.0% | 1.2% | 0.0% | 0.0% | 0.0% | 3.8% | 1.3% | 2.5% | 0.0% | 0.0% |
|  | Othereukaryotes | 4.0% | 3.4% | 0.0% | 0.0% | 0.6% | 6.6% | 3.4% | 2.2% | 0.0% | 0.0% |
|  | Bacteria | 0.0% | 0.2% | 0.9% | 0.9% | 0.0% | 0.0% | 0.0% | 0.0% | 0.0% | 0.0% |

